# Generation of adult hippocampal neural stem cells occurs in the early postnatal dentate gyrus and depends on cyclin D2

**DOI:** 10.1101/2022.12.05.518892

**Authors:** Oier Pastor-Alonso, Anum Syeda Zahra, Bente Kaske, Fernando García-Moreno, Felix Tetzlaff, Enno Bockelmann, Vanessa Grunwald, Soraya Martin-Suarez, Kristoffer Riecken, Otto Wilhelm Witte, Juan Manuel Encinas, Anja Urbach

## Abstract

In the hippocampus, lifelong neurogenesis is maintained by a pool of multipotent adult neural stem cells (aNSCs) residing in the subgranular zone of the dentate gyrus (DG). Yet, the mechanisms guiding the transition of NSCs from developmental to adult remain unclear. By using nestin-reporter mice deficient for D2, a cyclin expressed mainly postnatally, we show that the aNSC pool is established through D2-dependent proliferation during the first two weeks of life. The absence of D2 allows the normal development of the DG until birth but prevents the postnatal formation of radial glia-like aNSCs. Additionally, retroviral fate mapping demonstrates that aNSCs are born on-site from precursors located in the DG shortly after birth. Altogether, our data suggest that aNSCs are a population distinct from developmental NSCs and thus imply that adult hippocampal neurogenesis is not a mere continuation of development.

## INTRODUCTION

The DG is one of the few brain areas where neurogenesis persists into adulthood. The newly born neurons integrate and contribute to hippocampal functions including learning, memory and mood regulation, and alterations in their production have been associated to mental and neurological diseases (Toda et al., 2019). The source of this process lies in a pool of quiescent adult neural stem cells (aNSCs), which, upon activation, undergo mostly asymmetric division to self-renew and give rise to new neurons (Encinas et al., 2011; Kempermann, 2015). While symmetric NSC division is also possible (Bonaguidi et al. 2011), the division-coupled depletion of aNSCs outweighs their capacity for self-renewal, leading to a decline in neurogenesis with age (Encinas et al., 2011; Ibrayeva et al., 2021; Pilz et al., 2018). Consequently, the neurogenic capacity of the adult DG is largely determined by the initial size of the aNSC pool established during development.

The actual origin of aNSCs and the mechanisms controlling their establishment in the DG remain poorly understood. Adult NSCs reside in a special abventricular niche, the subgranular zone (SGZ), which is formed by postnatal day (P) 14 in mice (Nicola et al., 2015; Seki et al., 2014). Morphogenesis of the DG starts at late gestation and continues into postnatal periods (Altman and Bayer, 1990b; Nelson et al., 2020; Seki et al., 2014). A key feature of DG development is the formation of a dentate migratory stream (DMS) by developmental NSCs (dNSCs) and their progeny, which detach from the dentate neurepithelium (DNe) to form multiple transient niches and ultimately the DG (Altman and Bayer, 1990a; Li et al., 2009). Although dNSCs enter the nascent DG at embryonic stages, forming new hilar and subpial precursor niches inside the DG, the DMS remains active during the early postnatal period (Altman and Bayer, 1990a; Hodge et al., 2012).

Different studies suggest that aNSC precursors migrate along the DMS during embryonic development before colonizing the subgranular zone (SGZ), where they persist life-long (Berg et al., 2019; Hodge et al., 2013; Li et al., 2009). Others suggest that Sonic hedgehog-responsive precursors originating from the ventral hippocampus at E17.5 serve as a source of aNSCs (Li et al., 2013; Noguchi et al., 2019). Genetic fate mapping studies using HopxCreER^T2^ mice showed that aNSCs and dNSCs share a common embryonic origin and continuity in fate specification, suggesting that adult neurogenesis is a continuation of development (Berg et al., 2019). Although transcriptome analysis of the developmental and adult neurogenic cascade supported this perspective, the molecular profiles of aNSCs and dNSCs were significantly different (Berg et al., 2019; Hochgerner et al., 2018; Matsue et al., 2018; Valcarcel-Martin et al., 2020). Regardless of their embryonic origin, early postnatal cell division has been suggested to play a key role for the formation of aNSCs (Ortega-Martinez and Trejo, 2015; Youssef et al., 2018), supporting the idea that aNSCs could be an independent population generated during a critical postnatal period.

Previous work suggests that adult hippocampal neurogenesis requires cyclin D2 (D2), one of three D-cyclins essential for the progression of cells through the G1 restriction point in response to mitogens (Ansorg et al., 2012; Kowalczyk et al., 2004). We showed that this requirement builds up progressively during the early postnatal period, culminating in a proliferative arrest in D2-deficient mice between P14 and P28 (Ansorg et al., 2012). The fact that this period coincides with the formation of the aNSC pool and that the proliferative deficit of these mice cannot be overcome by exposure to neurogenic stimuli (Jedynak et al., 2012; Kowalczyk et al., 2004) led us to hypothesize that D2 might be relevant in the establishment of the aNSC pool. Here, we applied D2 knockout (D2KO) mice and targeted *in vivo* retroviral injections to describe when, how and where dNSC precursors divide and give rise to long-lived aNSCs. We found that D2-dependent proliferation is crucial for the postnatal formation of aNSCs and that the final mitosis generating aNSCs takes place on-site, in the DG during the first week and a half of life.

## RESULTS

### Inactivation of D2 depletes the population of self-renewing aNSCs

Previous studies suggest that D2 is critical for maintaining adult neurogenesis (Ansorg et al., 2012; Kowalczyk et al., 2004), but its function in aNSCs remains largely unclear. To specifically investigate the role of D2 in aNSCs, we used D2KO and WT littermates expressing GFP under control of the nestin promoter. In contrast to the densely populated WT SGZ, the SGZ of D2KO mice was virtually devoid of GFP^+^ cells with radial glia-like phenotype (Figure 1A). Most of the remaining GFP^+^ cells displayed a complex morphology. To quantify these differences in detail, we performed an unbiased 3D-Sholl analysis on randomly selected GFP^+^GFAP^+^ cells. While the Sholl profiles of WT NSCs reflected the prototypical morphotype with a single primary process and distal ramifications (Figure 1B), NSCs from D2KO mice displayed more proximal intersections and a smaller maximal radius (Figure 1B, C), confirming their higher complexity.

**Figure 1.**
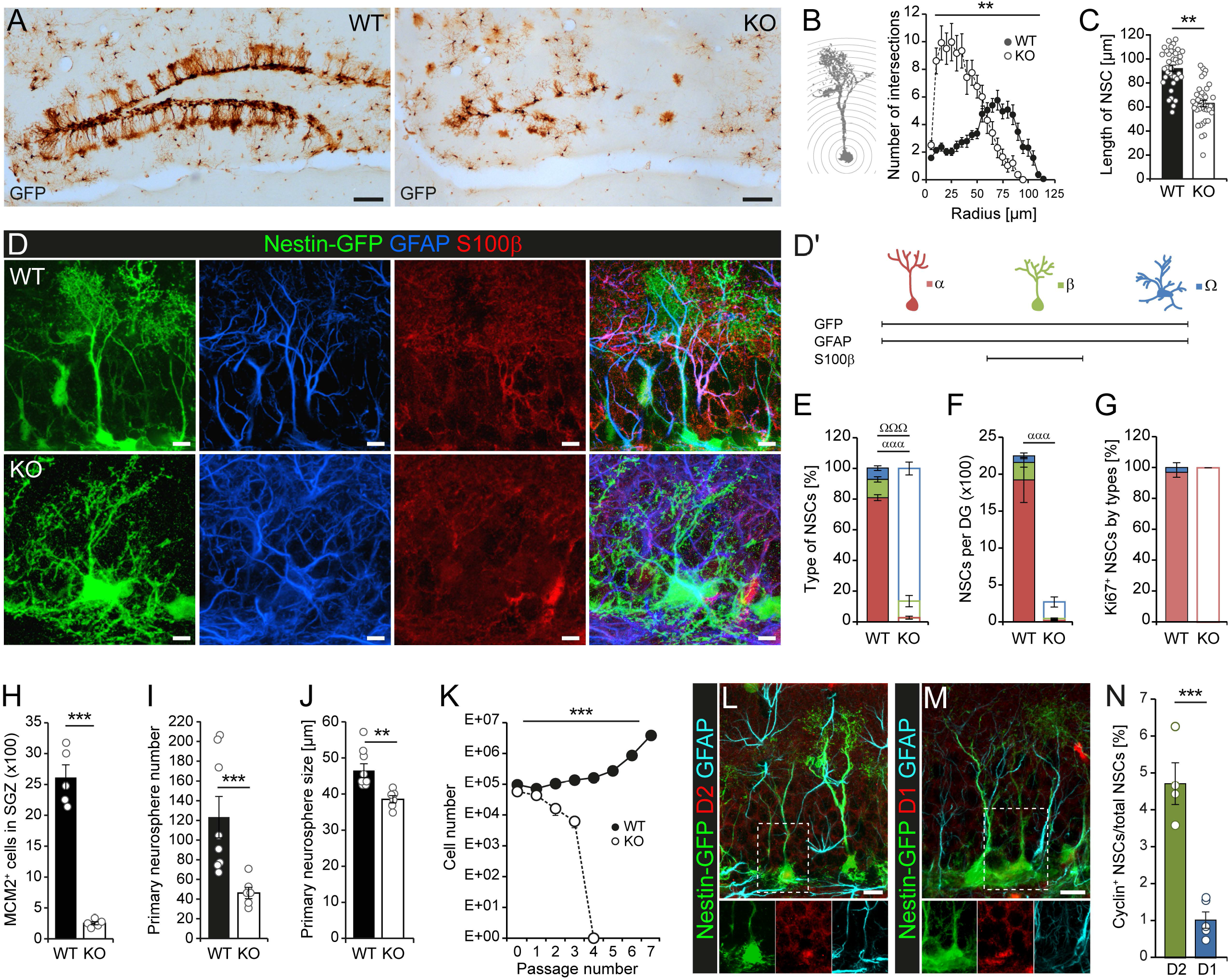
Lack of D2 results in a loss of NSCs in the adult DG. (A) ABC peroxidase staining of adult WT and D2KO nestin-GFP mice illustrating the loss of aNSCs in D2KO mice. Scale bar 100 μm. (B-C) 3D-Sholl analysis of randomly selected GFP^+^ NSCs (n = 4 mice/group, 8-10 cells/mouse) demonstrating increased complexity (B) and decreased length (longest axis with hilus-to-molecular-layer orientation; C) of mutant NSCs. (D) Confocal images of GFP, GFAP and S100β immunostained NSCs illustrating the phenotypical differences between prototypical NSCs in WT and NSCs of D2KO mice. Scale bar 10 μm. (E, F) Classification of NSC types based on the expression of S100β and morphological criteria showing the percentage (E) and total (F) changes of aNSC pool composition in D2KO mice. (G) Proportion of NSC types within the dividing (Ki67^+^) NSCs population. (H) Quantification of actively dividing (MCM2^+^) cells in the SGZ of D2KO and WT mice (n = 5/group). (I-J) Neurosphere-forming capacity of the DG during primary culture (n = 6-8/group). (K) Long-term self-renewal capacity of WT and D2KO cultures (n = 3-4/group). (L-N) Confocal images (L, M) and quantification (N) of D2^+^ and D1^+^ aNSCs (n = 4-5 /group). Scale bar 10 μm. Statistics: 2-way RM-ANOVA (B, E-G, K), Student’s t-test (C, J, N), Welch’s t-test (H), Mann-Whitney-U test (I); *p < 0.05, **p < 0.01, ***p < 0.001.

To determine whether the numerical and morphological changes in mutant NSCs reflect an imbalance in the composition of the NSC pool, we classified GFP^+^GFAP^+^ NSCs into three previously described subtypes (Figure 1D; Gebara et al., 2016; Martin-Suarez et al., 2019). Confirming these earlier reports, most NSCs of WT mice were radial glia-like α-cells (Figure 1E, F). The D2KO led to a disproportionate loss of α-cells (Figure 1E), that were almost absent in absolute terms (Figure 1F). Instead, most mutant NSCs were Ω-cells (Figure 1E, F), known to represent a deeply quiescent NSC subtype (Martin-Suarez et al., 2019). Thus, we co-stained against Ki67 and confirmed that α-cells represent the majority of dividing NSCs regardless of genotype, while Ω-cells rarely divide (Figure 1G). Accordingly, the SGZ of D2KO mice was virtually devoid of actively dividing cells (Ansorg et al., 2012; Figure 1H).

To corroborate these *in vivo* data, we performed neurosphere assays with serial passaging which help to address mitotic potential *in vitro.* Although neurospheres can be derived from aNSCs and their transit-amplifying progeny, only true NSCs are capable to self-renew over extended periods of time (Reynolds and Rietze, 2005; Walker and Kempermann, 2014). We found that cultures derived from young adult D2KO mice formed significantly less and smaller primary spheres than those of WT mice (Figures 1I, J), consistent with a numerical and functional deficit in aNSCs. When subcultured, mutant cells failed to expand and exhausted after a few passages (Figure 1K), confirming the lack of self-renewing aNSCs in the D2KO. Together, our *in vivo* and *in vitro* data demonstrate that the absence of D2 leads to selective loss of the radial glia-like, self-renewing population of aNSCs.

To further evaluate the role of D2 in the adult neurogenic lineage, we examined the expression of D2 in the adult WT SGZ (Figure S1A-G). Although all precursor types from aNSCs to neuroblasts were represented in the D2^+^ cell population, it mainly comprised transit-amplifying stages (Figure S1G). Moreover, D2 was found to be the predominant D-cyclin of radial glia-like NSCs (Figure 1L-N) compared with D1, the other D-cyclin expressed in the adult SGZ (Figure S1B-C, E). Co-staining with MCM2 revealed that the majority of D2^+^ cells were actively dividing, while the D1^+^ population was mostly quiescent (Figure S1H-J), altogether suggesting that D2 is required for lineage amplification during early stages of adult hippocampal neurogenesis.

### D2 is expressed in dNSCs

Although the results obtained in adult WT mice indicate that D2 is important for aNSC’s activation, the severity of the D2KO phenotype actually suggests a requirement for the formation of the aNSC pool. Cell division during early postnatal development has been suggested to play a role in the establishment of the aNSC population (Ortega-Martinez and Trejo, 2015; Youssef et al., 2018). Hence, we examined the expression of D-cyclins in radial glia-like NSCs at five postnatal stages (P0, P7, P10, P14 and P28), covering the period from when the DG is highly immature and NSCs are not considered adult (dNSCs; P0) to the age when NSCs with adult properties (aNSCs) occupy the SGZ (P14 onwards; (Berg et al., 2019; Nicola et al., 2015; Figures 2 and S2). We found that D2 is expressed by NSCs at all developmental stages, especially during the first 10 days of life when most of them were D2^+^ (Figure 2B-C). After a peak of expression at P7, the proportion of D2^+^ NSCs continuously declined, reaching residual adult levels at P28 (Figure 2C). Further quantification of D2^+^ and D2^-^ NSC numbers revealed that D2^+^ NSCs are a transient population that appears shortly after birth and increases significantly until P10, before declining and returning to nearly P0 levels at P28 (Figure 2D). Next, we examined D1, which is also expressed by aNSCs (Figure 1 M-N) but appears largely dispensable for adult neurogenesis (Kowalczyk et al., 2004). Except for P0, when more NSCs expressed D1 than D2 (Figure 2C), the proportions and absolute numbers of D1^+^ NSCs followed a similar time course as those expressing D2 (Figures 2C, E and S2). The high expression of both D-cyclins in dNSCs indicates a substantial overlap of expression. Therefore, we calculated a minimum coexpression index for each age and determined the zero of the curve fit. This approach revealed that D2 and D1 must be co-expressed in a fraction of dNSCs until at least P13 (Figure 2F).

**Figure 2.**
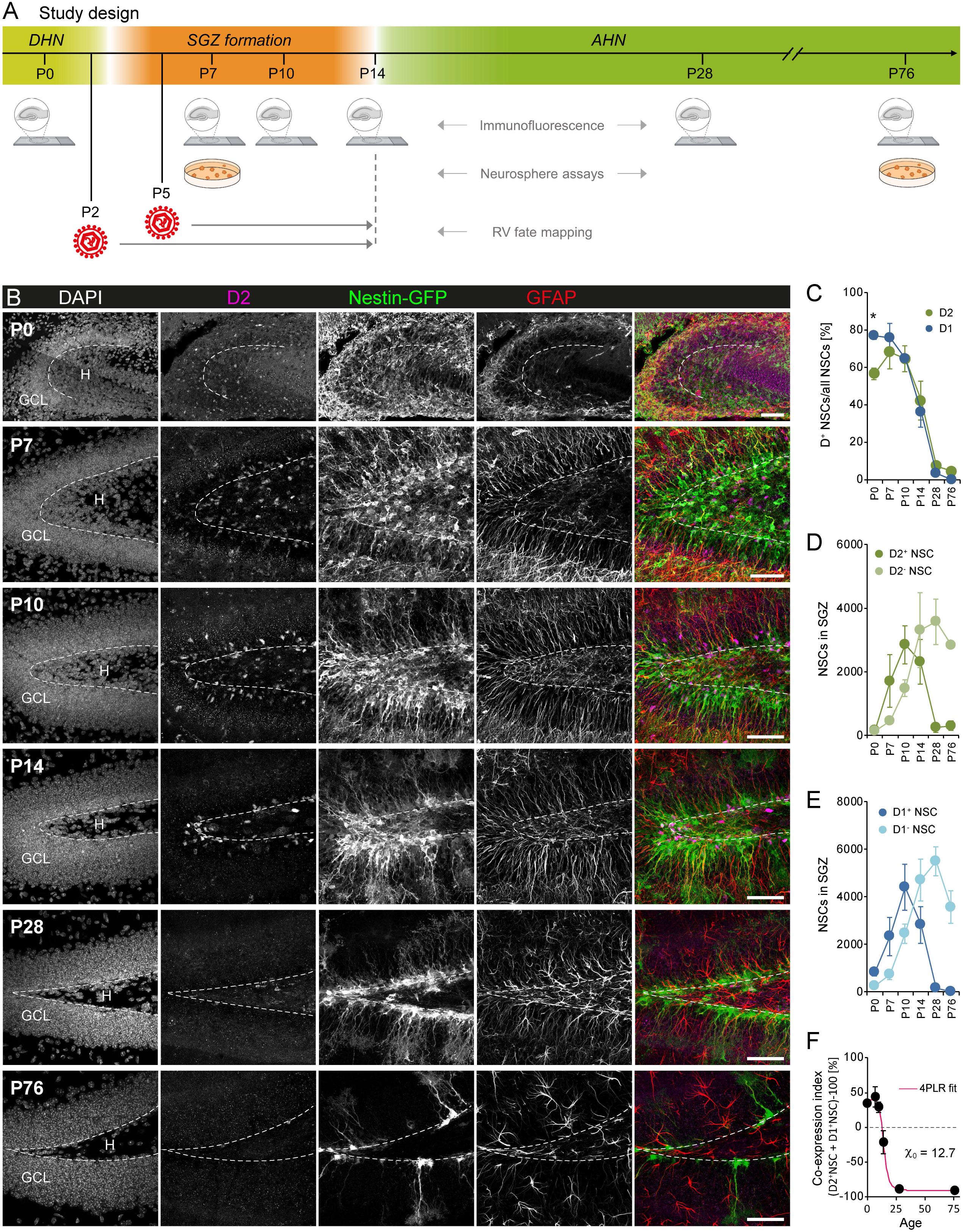
D2 as well as D1 are expressed in NSCs of the postnatal DG. (A) Schematic of the study design for investigating the SGZ niche in WT and D2KO mice. (B) Confocal images illustrating the expression of D2 in the hilar and SGZ niche of the developing DG. Scale bars 50 μm. GCL, granule cell layer; H, hilus. (C) Proportions of NSCs expressing D2 or D1 (n = 4-5/group). See also Figure S2. (D) Total number of D2^+^ and D2^-^ NSCs (n = 4/group). (E) Total number of D1^+^ and D1^-^ NSCs (n = 4-5/group). See also Figure S2. (F) Coexpression index of D2 and D1 in NSCs to estimate how long their expression must overlap in NSCs. Curve fit represents a Four Parameter Logistic (4PL) Regression. Statistics: 2-way ANOVA, *p < 0.05.

In parallel to the D-cyclin^+^ NSCs, but with a slight delay, the populations of D^-^ NSCs increased from P7 and continued expanding until P14, after P28 they began to decline (Figure 2 D, E). These data demonstrate that both D-cyclins are highly expressed by dNSCs during the time of peak proliferation in the developing DG (Altman and Bayer, 1990a; Bond et al., 2020) and are then downregulated making the NSCs transition to a quiescent adult state (Berg et al., 2019).

### D2 deficiency impairs the formation of the aNSC pool

Next, we examined the development of the aNSC pool in D2KO mice. We previously showed that overall proliferation is reduced in the SGZ of D2KO mice as early as P7 (Ansorg et al., 2012). To evaluate whether the D2KO alters the cycling activity of dNSCs, we co-stained them with Ki67 (Figure 3A). Interestingly, the proportion of dividing NSCs in the D2KO was unaltered (Figure 3B). However, in total numbers, significantly fewer Ki67^+^ NSCs were found in the DG of D2KO mice from P14 onwards (Figure 3C), suggesting that the number of NSCs produced, rather than their propensity to divide, was decreased due to the absence of D2. Accordingly, quantification of total NSCs in the SGZ revealed a progressive deficit in aNSC pool amplification from P7 onwards (Figure 3D). Consistent with other studies (Glickstein et al., 2007a; Kowalczyk et al., 2004), the overall architecture of the mutant DG at perinatal age was comparable to that of WT mice. Moreover, at P0, the number of dNSCs was similar in both groups, indicating that embryonic development of the DG and its germinative niches is not significantly affected by the deletion of D2. From P0 to P7, the numbers of NSCs increased in both WT and D2KO mice, although tending to be higher in WT mice. Between P7-P10, when NSCs transition from developmental to adult, WT mice displayed the greatest NSC population growth (Figure 3 D-E) which failed to occur in D2KO mice (Figure 3D). After P10, NSCs of KO mice gradually disappeared, increasing the difference to WT further (Figure 3D). From P14 onwards, a decline in NSC numbers was noticeable also in WT mice (Figure 3A, D), suggesting that the balance between aNSC generation and depletion had already tipped in favor of exhaustion. Further investigation of the mutant NSC pool revealed that the transient D1^+^ dNSC population, albeit in smaller numbers, is formed also in D2KO mice (Figure S2B, D), whereas the expansion of the D1^-^ population is completely prevented (Figure S2C). This suggests that D1 contributes to dNSC proliferation but cannot compensate for the lack of D2 during aNSC amplification. The impaired formation of the aNSC pool observed in mutant mice was accompanied by a reduced growth of the SGZ, resulting in a significantly smaller SGZ from P10 onwards (Figure 3F). These data suggest that D2 becomes increasingly required for DG neurogenesis during the first two weeks of life, whereas it is largely dispensable for early dNSC proliferation. This was confirmed in neurosphere assays with NSCs isolated from the P7 DG (Figure 3G-I). Even though cultures of D2KO mice produced fewer primary neurospheres (Figure 3G), their size was similar (Figure 3H) and they were able to grow exponentially (Figure 3I), demonstrating the presence of self-renewing NSCs in the early postnatal DG of mutant mice.

**Figure 3.**
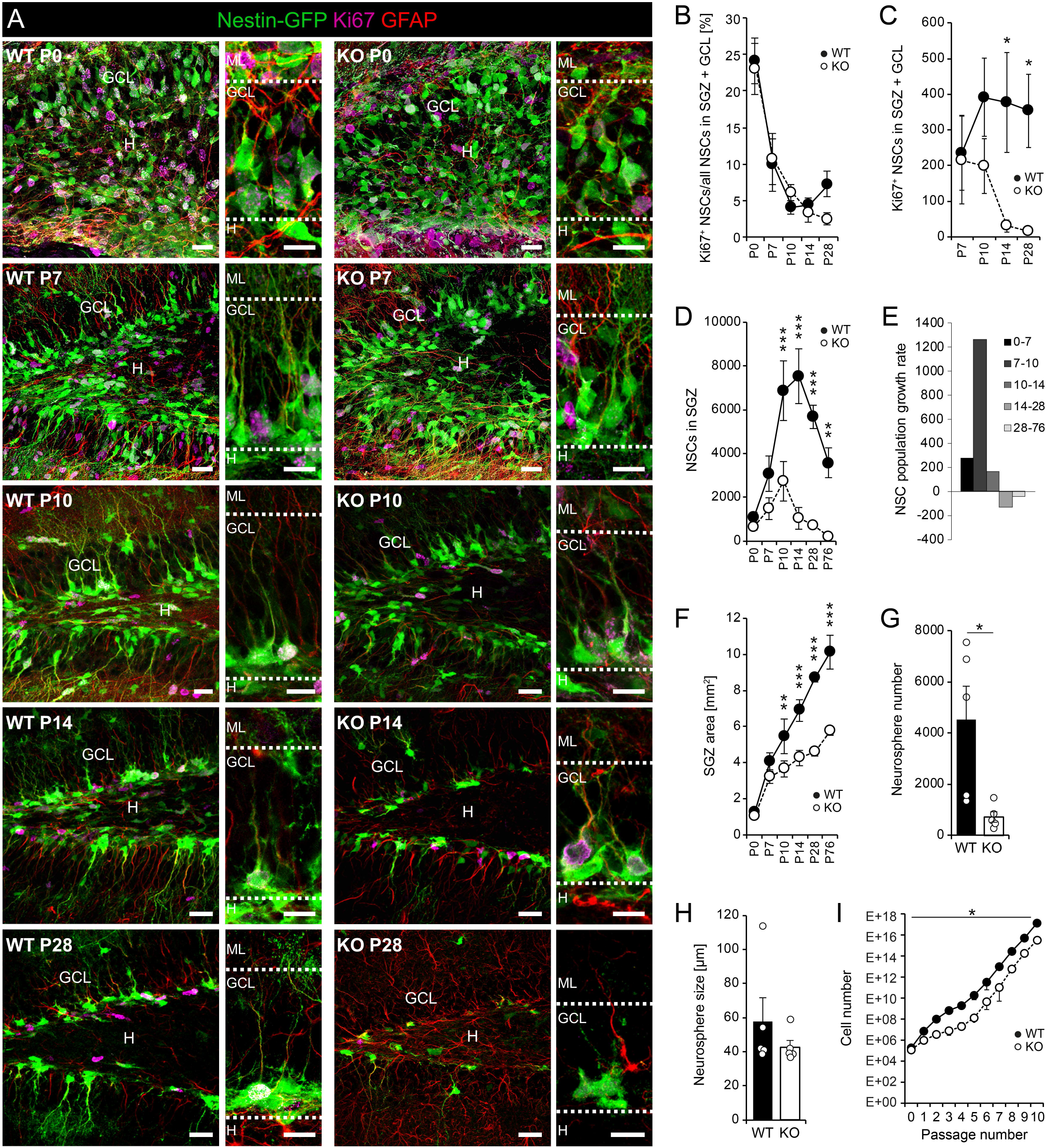
Absence of D2 impairs the generation of aNSCs. (A) Confocal images of nestin-GFP, Ki67 and GFAP expression in the developing DG of WT and D2KO mice. Scale bars 25 μm in lower magnification images, 10 μm in higher magnification images. (B-C) Percentage (B) and absolute number (C) of proliferating (Ki67^+^) NSCs. (D) NSC quantification showing that the D2KO prevents the expansion of the aNSC population during the first 2 weeks of life (n = 4-5/group). (E) Average NSC pool growth rate per day in WT, demonstrating the strongest expansion between P7 and P10. (F) Quantification of the SGZ area (n = 4-5/group). (G-H) Primary neurosphere-forming capacity of postnatal DG precursor cells isolated at P7 (n = 5/group). (I) Long-term self-renewal capacity of P7 cultures (n = 3/group). Statistics: 2-way ANOVA (B-D, F), Welch’s t-test (G), Mann-Whitney-U test (H), 2-way RM-ANOVA (I); *p < 0.05, **p < 0.01, ***p < 0.001. Horizontal bars in (C) and (I) represent a main effect of genotype.

To examine whether the lack of aNSCs in the mutant DG was indeed caused by a deficit in their generation instead of premature differentiation or death of dNSCs, we quantified the numbers of astrocytes, neuroblasts and apoptotic cells in the different layers of the developing DG (Figure S3). GFAP^+^ stellate astrocytes of different developmental stages (immature: GFP^+^, mature: GFP^-^) were found to a similar extent in both groups, except for the postnatal germinative niches in the subpial zone and the hilus of KO mice, which contained fewer astrocytes in comparison to WT mice (Figure S3A). The number of apoptotic cells in the mutant SGZ was actually smaller from P7 onwards, consistent with fewer NSCs and reduced neurogenesis, but similar to WT in the hilus (Figure S3B). As well, numbers of DCX^+^ neuroblasts in the SGZ (Figure S3C) and the volume of the GCL were smaller in KO mice (Figure S3D). The finding that neither neurogenesis, astrogliogenesis nor cell death are increased during DG development of mutant mice supports the assertion that the D2KO directly impairs the generation of aNSCs.

Taken together, these data demonstrate the requirement of D2 for the late-stage cell divisions that lead to the formation of aNSCs in the early postnatal DG. Even though deficiency of D2 does not affect regular ontogenesis of the DG during the fetal period, it prevents the generation of aNSCs, resulting in premature loss of neurogenic capacity in the juvenile DG.

### Spatially restricted *in vivo* retroviral labeling reveals the postnatal origin of aNSCs

Next, we determined where exactly aNSC precursors divide and give rise to aNSCs. Through stereotactic SFFV-RV-mCherry infections into nestin-GFP pups (Mignone et al., 2004), we specifically compared the contribution of precursors migrating along the DMS to those already located inside the postnatal DG (Altman and Bayer, 1990a; Berg et al., 2019; Hodge et al., 2013; Sugiyama et al., 2013). After validating the spatial precision of the retroviral injections into the DG and the DMS (Figure 4C, Figure S4A-B), we performed injections at P2 or P5, representing ages before and close to the peak generation of aNSCs (Berg et al., 2019; Ortega-Martinez and Trejo, 2015), and sacrificed the mice at P14, when the aNSCs pool is fully established. In all cases, mCherry^+^ neurons with prototypical dendritic arborization extending into the ML and axons extending into the hilus were found (Figure 4A-B; Figure S4C-D). Strikingly, mCherry^+^ aNSCs were only found when RV-SFFV-mCherry was injected into the DG but not into the DMS (Figure 4D-E). This result indicates that aNSCs are generated exclusively from precursors located in the DG early after birth, while precursors dividing in the postnatal DMS do not contribute to the formation of aNSCs and adult neurogenesis. Summarizing these results, we conclude that the persistent aNSC population is generated on-site in the DG via division of precursor cells that colonized the DG before P2.

**Figure 4.**
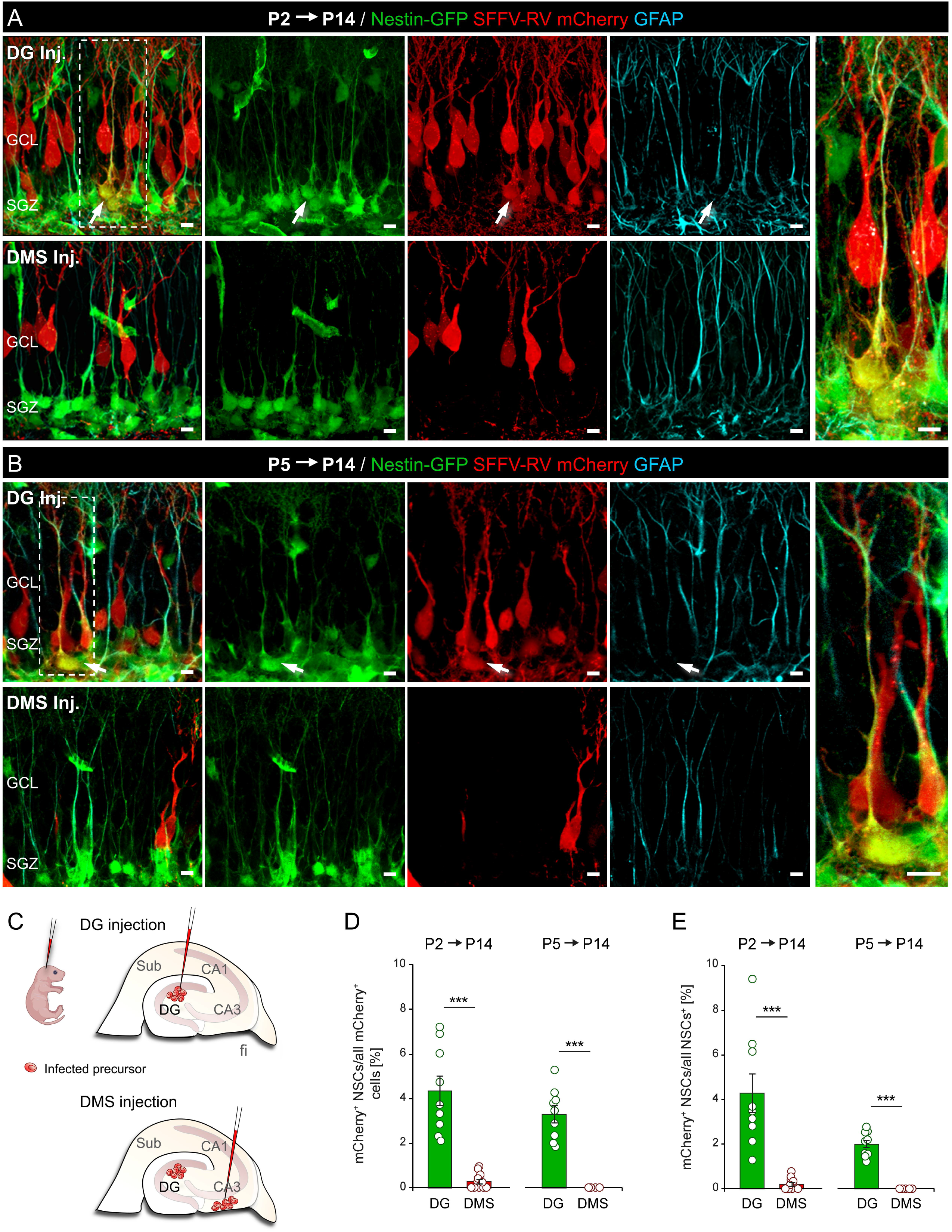
Adult NSCs are generated on-site in the early postnatal DG. (A-B) Confocal images of the nestin-GFP/GFAP-immunostained DG of P14 mice after SFFV-RV mCherry injection into either the DG or the DMS at P2 (A) or P5 (B). DMS, dentate migratory stream. Scale bars 5 μm. (C) Schematic representation of the DG and DMS retroviral injections. (D) Percentage of NSCs among all infected (mCherry^+^) cells (n = 5-12 mice/condition). (E) Percentage of mCherry^+^ NSCs among all NSCs (n = 5-12 mice/condition). Statistics: Welsh’s t-test (D, E) and Student’s t-test (E), ***p < 0.001.

## DISCUSSION

Our study reveals the indispensable role of D2 in the generation of the hippocampal aNSC pool. The D2-driven formation of aNSCs during a discrete time window (P0-P14) from specific precursors that enter the DG perinatally strongly suggests that aNSCs and adult neurogenesis are distinct from, and not simply a continuation of, those during hippocampal development.

Although D-cyclins are interchangeable in many cellular contexts (Sherr and Roberts, 2004), several lines of evidence suggest that they possess unique roles during nervous system development (Glickstein et al., 2007a; Glickstein et al., 2007b; Lukaszewicz and Anderson, 2011; Wianny et al., 1998). In the DG, deletion of D2 but not of D1 leads to a severe deficit in neurogenesis that builds up progressively during the early postnatal period (Ansorg et al., 2012; Kowalczyk et al., 2004). Because this period coincides with the appearance of aNSCs in the SGZ (Nicola et al., 2015; Ortega-Martinez and Trejo, 2015), we hypothesized that the impaired neurogenesis of D2KO mice was due to the failed generation of the population of aNSCs during DG development. Our results show that deletion of D2 indeed prevents the formation of the radial glia-like NSC pool early on and that the few NSCs still born deplete much earlier than usual. This effect could not be compensated by D1, which, however, is sufficient for assuring the prenatal development of the DG and the proliferation of dNSCs in the perinatal DG. The cause for the differential requirement for D-cyclins in dNSCs and aNSCs has not been established, but may involve the preferential regulation of D1 and D2 by spatiotemporally changing niche signals and differences in their non-canonical effects on downstream molecular pathways and gene expression (Hydbring et al., 2016; Pagano and Jackson, 2004). In this context, it could be speculated that D2 acts as an indispensable effector of sonic hedgehog signaling, which is required for the establishment and expansion of aNSCs and whose inactivation leads to a similar, but more severe, phenotype than the D2KO (Han et al., 2008; Li et al., 2013; Noguchi et al., 2019). How exactly D2 controls the formation of aNSCs remains to be determined. Previous reports indicate that D-cyclins, in addition to their role in proliferation, are also involved in fate specification and survival of various types of stem cells (Choi et al., 2014; Glickstein et al., 2007a; Glickstein et al., 2009; Lukaszewicz and Anderson, 2011; Pauklin et al., 2016). However, we did not observe signs of increased cell death, neuron or astrocyte production in the early postnatal niches of mutant mice, indicating that D2 directs the cell divisions leading to the formation of aNSCs. Interestingly, the proportion of actively dividing NSCs in the SGZ of postnatal D2KO mice was unchanged, arguing against a general proliferation defect. This combined with the concomitant deficit in aNSC pool expansion suggests a requirement for D2 in fate specification during divisions producing aNSC, similar to what has been observed in the embryonic cortex (Glickstein et al., 2007a; Glickstein et al., 2009; Tsunekawa et al., 2012). Moreover, it is conceivable that D2 specifically promotes symmetric NSC divisions required for the expansion of the aNSC pool, whereas D1 drives asymmetric divisions of dNSCs that foster developmental neurogenesis, consistent with a model proposed for embryonic radial glia cells (Glickstein et al., 2009). To precisely determine the role of D2 in aNSC development, it will be necessary to generate mice that allow conditional deletion of D2 from NSCs and the fate mapping of D2-expressing precursor cells.

The spatial origin of aNSCs remains a controversial topic. Whereas precursors from the DNe migrating along the DMS have traditionally been considered the source of aNSCs (Berg et al., 2019; Hodge et al., 2013; Li et al., 2009), others suggest the ventral sector of the hippocampus as origin of aNSCs (Li et al., 2013; Noguchi et al., 2019). Regardless of the embryonic origin, we aimed to determine where the final, D2-dependent mitoses that give rise to persistent aNSCs take place. Previous retroviral lineage tracing experiments in rats suggest that precursors residing in the hilar germinative matrix serve as a source of aNSCs (Namba et al., 2005). Moreover, studies in which pCAG-GFP was electroporated in the ventricular zone of mice imply that NSCs exiting the DNe later than E16 only generate neurons (Ito et al., 2014). Our fate mapping analyses show that aNSCs originate from precursor divisions inside the early postnatal DG, while precursors dividing at the same time in the DMS are purely neurogenic. Thus, besides uncovering a mechanism by which aNSCs are generated in the postnatal DG (D2-dependent mitosis), we show that aNSC precursors enter the DG before P2 and form a population separate from the exclusively neurogenic precursors that continue to migrate from the DNe into the postnatal DG.

These results challenge previous ideas that conceive aNSCs as remnants of embryonic DG development. In spite of common features with dNSCs, aNSCs have different dynamics of cell division, shift their mode of regulation from intrinsic to extrinsic and differ in their transcriptomic landscape and marker expression (Borrett et al., 2022; Hochgerner et al., 2018; Matsue et al., 2018; Valcarcel-Martin et al., 2020). All this strongly suggests that adult hippocampal neurogenesis is a different process from developmental neurogenesis, which is supported by recent work showing that the maturation and morphological properties of adult-born neurons differ from those that are neonatally born (Cole et al., 2020; Kerloch et al., 2019). Our results are not at odds with aNSCs being an endpoint of one developmental lineage (Berg et al., 2019). However, they demonstrate that aNSCs represent a separate population that is produced and silenced postnatally to serve as a neurogenic reserve for life-long neurogenesis. This is consistent with the set-aside model proposed for V-SVZ NSCs (Fuentealba et al., 2015; Furutachi et al., 2015), yet on a different time scale.

Our work solves the long-standing question about the formation of hippocampal aNSCs and identifies the critical period and spatial location in which they are born. Symmetric division (Bonaguidi et al., 2011) as well as time-dependent changes in cell division and self-renewal (Bottes et al., 2021; Harris et al., 2021; Ibrayeva et al., 2021) are able to extend the productive life-span of the aNSC population. Still, the vast majority of the neurogenic output depends directly on the initial size of the aNSC population. An important prediction of these and previous findings (Ortega-Martinez and Trejo, 2015; Youssef et al., 2018) is that any pathophysiological event interfering with the establishment of aNSCs during this critical period will lead to lasting impairments of adult hippocampal neurogenesis and increased susceptibility to pathology.

The discovery of how and where aNSCs are generated in the rodent DG opens avenues for their manipulation and understanding of their cell nature in both rodents and humans. The existence of D2-dependent division leading to the formation of long-lasting precursors in the human DG remains unsolved. Further research is required to understand the progression of dNSCs in the human DG and whether the process is comparable to rodent DG development.

## EXPERIMENTAL PROCEDURES

Detailed methods are provided in the online version of this paper.

## SUPPLEMENTAL INFORMATION

Supplemental Information can be found with this article at …

## ACKNOWLEDGMENTS

We thank the staff at the Leioa animal facility of UPV/EHU, Laura Escobar at the Imaging Core Facility in Achucarro, and the technicians in the Jena Neurology department for technical support. We would also like to thank members of the Neural Stem Cell and Neurogenesis group in Leioa and of the Hippocampal Plasticity and Neurogenesis group in Jena for discussion and insight. O.P.-A. received a UPV/EHU predoctoral fellowship. J.M.E. was supported by the Spanish Ministry of Economy and Competitiveness (MINECO; grants SAF2-015-70866-R, MINECO Ramón y Cajal Program: RYC 2012-11137 and MINECO PCIN-2016-128 (ERA-NET-NEURON III program)). O.W.W. received support from German Research Foundation (DFG; grants WI 830/12-1 and WI 830/12-2) and Marie Skłodowska-Curie Innovative Training Network (ITN; grant 859890 SmartAge). A.U. was supported by the Interdisciplinary Center for Clinical Research (IZKF; grant AMSP06). Some figures include icons from BioRender.com.

## AUTHOR CONTRIBUTIONS

Conceptualization, O.P.-A., A.S.Z., F.G.-M., J.M.E., A.U.; Methodology, O.P.-A., A.S.Z., F.G.-M., E.B., A.U.; Validation: O.P.-A., A.S.Z., A.U.; Formal Analysis, O.P.-A., A.S.Z., E.B., A.U.; Investigation, O.P.-A., A.S.Z., F.G.-M., B.K., E.B., F.T., V.G., J.M.E, A.U.; Resources, C.R., O.W.W., J.M.E., A.U.; Data Curation, O.P.-A., A.S.Z., F.G.-M., J.M.E., A.U.; Writing – Original Draft, O.P.-A., A.U.; Writing – Review and Editing, O.P.-A., A.S.Z., J.M.E., A.U.; Visualization, O.P.-A., J.M.E., AU.; Funding Acquisition, O.W.W., J.M.E., A.U.; Supervision, F.G-M., O.W.W., J.M.E., A.U.

## SUPPLEMENTAL FIGURE LEGENDS

**Figure S1 (related to Figure 1). D2 is expressed by actively dividing progenitors in the adult SGZ.** (A, B) Peroxidase staining of coronal brain sections illustrating the distribution of D2 (A) and D1 (B) in the DG of WT and D2KO mice. Upper pannels: overview of the DG, lower pannels: magnified image of the suprapyramidal blade. (C) Density of cyclin D^+^ cells in the adult SGZ. (D-F) Expression of *ccnd1* and *ccnd2* mRNA and lack of *ccnd3* in the adult DG. Allen Mouse Brain Atlas, https://mouse.brain-map.org/experiment/show/205, https://mouse.brain-map.org/experiment/show/69540507, https://mouse.brain-map.org/experiment/show/68191468. (D’-F’) Expression mask image display highlighting cells with highest probability of gene expression. (G) Quantification of the proportions of NSCs, type 2 cells and neuroblasts among D2^+^ cells revealed a prevalence of transit-amplifying type 2 progenitors. (H, I) Confocal images illustrating the expression of MCM2 in D2^+^ and D1^+^ cell populations. (J) Proportions of D2^+^ and D1^+^ cells expressing MCM2. Data represent mean ± SEM, n ≥ 4 per group; statistics: One-way ANOVA (C, J) and One-way RM-ANOVA (G), **p < 0.01, ***p < 0.001; Scale bars: 25 μm (A-B, H-I) and 200 μm (D-F’). Abbreviations: ML, molecular layer; GCL, granule cell layer; H, hilus.

**Figure S2 (related to Figure 2). The lack of D2 impairs the postnatal expansion of the D1^-^ NSC population and diminishes the transient D1^+^ NSC population.** (A) Confocal images of the developing DG of WT and D2KO mice immunostained against D1, nestin-GFP, GFAP and DAPI. Figures represent maximum intensity projections of 4.3 μm (P0) and 13.5 μm (P7 to P76) high z-stacks. (B, C) Quantification of D1^+^ and D1^-^ NSC numbers in the SGZ. (B) D1^+^ NSCs are a transient population. Deletion of D2 affects but does not prevent their appearance, suggesting that D1 may either partially compensate for the lack of D2 or designate a distinct dNSC population. (C) Deletion of D2 prevents the expansion of the D1^-^ NSC population. (D) Proportions of NSCs expressing D1 are higher in mutant mice compared to WT. Data represent mean ± SEM, n = 4-5 per group; statistics: 2-way ANOVA, **p < 0.01, ***p < 0.001; Scale bars: 50 μm. Abbreviations: H, hilus; GCL, granule cell layer.

**Figure S3 (related to Figure 3). The impaired formation of the aNSC pool in D2KO mice is not caused by apoptosis or differentiation of postnatal NSCs.** (A) Postnatal astrogliogenesis is not increased upon deletion of D2. Top panel (from left to right): Sample confocal images of immature (GFAP^+^GFP^+^) and mature (GFAP^+^GFP^-^) stellate astrocytes in the ML and of an immature polar astrocyte in the GCL. Images represent maximum intensity projections of 12.07 μm confocal Z-stacks; scale bars 20 μm and 10 μm (magnified images). (A’) Quantification of immature and mature astrocytes. Because of the extensive migration of dNSC-derived astrocytes in the postnatal DG (Brunne et al., 2010), analysis was performed in different layers of the DG. (B) Quantification of apoptotic cells. Top panel: DAPI staining showing the morphotypes of nuclei (pyknotic, donut-shaped and karyorrhectic; single optical planes) considered as apoptotic. Scale bar 10 μm. (C, C’) Quantification of neuroblasts and immature neurons in the postnatal SGZ. Images represent maximum intensity projections of 6.9 μm confocal Z-stacks, scale bars represent 20 μm. (D) Quantification of the GCL volume (2-way ANOVA, ***p < 0.001). Values represent mean ± SEM, n = 4-5 mice/group. Abbreviations: SGZ, subgranular zone; GCL, granule cell layer; iML, inner half of the molecular layer; oML, outer half of the ML representing the subpial germinative niche of the developing DG.

**Figure S4 (related to Figure 4). Fluorescence slide-scanner images showing the spatial destination of SFFV-RV-mCherry-infected cells after injection into the DG or in the DMS.** (A, B) To evaluate the spatial precision of SFFV-RV infections into the DG or the DMS, C57Bl/6 mice were injected either at P2 (A) or P5 (B) and sacrificed 2 days later. The squares delimitate the DMS shown at higher magnification below, and arrows mark the injection path. At both ages, mCherry^+^ cells were observed exclusively at the infected site, verifying that the injection paradigm is effective for tracing the lineage of aNSCs from different postnatal niches. A wider distribution, including the DG near the fimbrodentate junction, was found in mice injected in the DMS, reflecting the migratory activity of the precursors located in that area. (C, D) Nestin-GFP expressing pups were injected at P2 (C) or P5 (D) and the spatial destination of infected cells from rostral to caudal was assessed at P14. The arrows mark the injection path. Insets delineate the DG, which is shown at higher magnification in C’ and D’. In all cases, we observed mCherry^+^ cells located in the GCL, as well as axons entering the hilus. However, mCherry^+^ cells connecting the LV with the DG were observable only when the injection was done in the DMS. Scale bars = 200 μm. Abbreviations: CA, Cornu ammonis; DG, Dentate gyrus; DMS, Dentate migratory stream; fi, fimbria; GCL, Granule cell layer; LV, Lateral ventricle.

## STAR METHODS

Detailed methods are provided in the online version of this paper and include the following:

### RESOURCE AVAILABILITY

#### Lead contact

Further information and requests for resources and reagents should be directed to and will be fulfilled by the corresponding authors, Anja Urbach and Juan Manuel Encinas

#### Materials availability

This study did not generate new reagents.

#### Data and Code Availability

This study did not generate/analyze datasets/code.

### EXPERIMENTAL MODEL AND SUBJECT DETAILS

**Animals**

### METHOD DETAILS

**Injection of retroviral vectors**

**Tissue preparation and immunostaining**

**Neurosphere assay**

### QUANTIFICATION AND STATISTICAL ANALYSIS

**Image capture and analysis**

**Statistical analysis**

## EXPERIMENTAL MODEL AND SUBJECT DETAILS

### Animals

All studies were carried out on cyclin D2 knockout/Nestin-GFP (D2KO) and wildtype/Nestin-GFP (WT) mice of mixed gender. The double-transgenic line was established by crossing heterozygous cyclin D2 knockout mice (Sicinski et al., 1996) with Nestin-GFP mice expressing enhanced green fluorescent protein (GFP) under control of rat nestin gene regulatory elements (Yamaguchi et al., 2000). The heterozygous offspring of this breeding was mated to each other to obtain Nestin-GFP expressing animals homozygous for the mutated (D2KO) or the wildtype (WT) *ccnD2* allele. All mouse lines were maintained on a C57BL/6 background. Mice were kept under specific pathogen free conditions on a 14 hours light/10 hours dark cycle with food and water ad libitum. Animals were weaned at around P30, hence mice of all groups until P28 were housed as litters and/or with their mothers. Adult mice were group-housed. To minimize litter effects, littermates were assigned to groups in such a way that mice from at least three different litters contributed to a particular group, despite in the P7 WT group, which was recruited from only two litters. All animal procedures were in strict compliance with the European animal welfare regulations (EU directive 2010/63/EU and 2007/526/EC guidelines) and approved by the local authorities (Thueringer Landesamt für Verbraucherschutz, Bad Langensalza, Germany; University of the Basque Country (EHU/UPV) and the Comunidad Foral de Bizkaia Ethics Committees).

### Cell Culture-Neurosphere assay

DGs of P7 and of 6-9 weeks old D2KO and WT mice were dissected and individually subjected to enzymatic and mechanical dissociation (Neural Tissue Dissociation Kit P, Miltenyi Biotec) as described earlier (Walker and Kempermann, 2014). The obtained single cells were resuspended in 20 ml serum-free Neurobasal media (Gibco) containing 2% B27 (Gibco), 1x Glutamax (Gibco), 20 ng/ml EGF (PeproTech), 20 ng/ml bFGF (PeproTech), 50 U/ml Pen/Strep (Gibco), 20 μg/ml heparin (Sigma), and seeded into 96-well plates (200 μl/well, resulting densities: 2-10 cells/μl). Primary cultures were maintained at 5% CO2 and 37°C for 10 days. The total number and size of spheres that formed in each well were determined microscopically at 200x magnification or with an IncuCyteZoom system (EssenBioscience, Germany). Only spheres larger than 20 μm in diameter were analyzed. To determine self-renewal capacity, we performed bulk culture serial passaging (10x 7 days) as described earlier (Walker and Kempermann, 2014). Neurospheres of each animal were pooled, digested with Accutase (Sigma) and triturated into single cell suspensions. After counting cell numbers using a hemocytometer, cells were reseeded at a density of 1 x 10^4^ cells/cm^2^. The fold expansion relative to the seeded cell number was calculated and multiplied with the total of the previous passage to determine the theoretical total cell number.

## METHOD DETAILS

### Injection of retroviral vectors

We adapted a method to perform intrahippocampal injections in accordance with our requirements: 1) the need to inject independently in two different regions (DG and DMS) of the hippocampus, and 2) its adaptation to the varying size of these regions during early postnatal development.

We traced the lineage of postnatal hippocampal progenitors with mCherry-expressing spleen-focus forming virus gamma-retroviruses (SFFV-RV; Gomez-Nicola et al., 2014). P2 and P5 old mice were anaesthetized by hypothermia by placing them in ice for 2 minutes and then secured to a platform placed in the stereotaxic apparatus. Aided by a fiber optic light, Lambda was localized by head transillumination and used as reference for the injections with a heat-pulled glass microcapillary at the following stereotaxic coordinates: for DG, −1 mm anteroposterior (AP), ±1.2 mm laterolateral (LL), −1.7 mm dorsoventral (DV); for the DMS −0.9 mm AP, ±1.4 mm LL, −1.7 mm DV. The microcapillary was introduced percutaneously to inject 0.3μl of SFFV-RV in each hippocampal region at a rate flow of 0.3μl/min. After surgery, mice were quickly placed in a warm water bath and thereafter kept on a thermal blanket until their respiration, skin color and locomotor activity returned to normal. Finally, to facilitate re-acceptance of the mother, they were impregnated with a mixture of the mother’s own stool, bedding and water prior to the returning to the breeding cage. The whole procedure for each pup was carried out in less than 10 minutes to minimize as much as possible maternal stress. No pups were rejected or abused by the dam. However, and despite mortality was almost negligible, all the pups that did not recover from anesthesia corresponded to the P5 age group, suggesting a major susceptibility to hypothermia at this age.

The implemented method allows the delivery of viral particles in the targeted anatomical regions, yet the injection accuracy is not 100% due to the variability in animal size and methodological limitations. Therefore, given the proximity of both injection sites and their small size at these postnatal stages, we employed *a posteriori* criteria to correctly assign injected animals to each group. To classify an injection as “DG”, we ensured that no cells were labeled by the RV in the DMS. Furthermore, the path of the microcapillary was observable in several cases, assuring the identification of the injection site. To classify injections in the “DMS” group, infected cells were required to populate the entire stream above the fimbria from the injection site into the DG. Injections that only reached the meninges could be distinguished from DMS injections due to their location forming a stream under the ventral blade of the DG and the characteristic elongated morphology of the cells in this region.

### Immunostaining

Brains were obtained from different postnatal/juvenile ages (P0, P7, P10, P14, P28; at least 4 mice per group) as well as from adult mice (approx. P76; at least 4 mice per group; Fig. 2A). The P0 pups were decapitated and their brains fixed in 2% paraformaldehyde (PFA; w/v) in 0.1 M phosphate buffer (pH 7.4) at 4°C for 20 hours. All other mice were sacrificed by an overdose of isoflurane and transcardially perfused with ice-cold PBS (pH 7.4; 5 ml/min for 2 min) followed by 4% PFA in 0.1 M phosphate buffer, (pH 7.4; 5 ml/min for 8 min). After dissection, brains were removed and post-fixed in PFA for 3 h at 4°C. For storage at −80°C, the brains were cryoprotected consecutively in 10% and 30% sucrose (in PBS, 4°C), cut midsagitally into the two hemispheres, frozen in 2-methylbutan (−25 to −30°C) and stored at −80°C. Hemispheres were sectioned either into 40 μm coronal sections on a sliding microtome (Epredia 400, Fisher Scientific, Germany) or into 50 μm sagittal sections,on a Leica VT 1200S vibratome (Leica Microsystems, Germany), and stored in antifreeze solution. For each antibody combination, every 6^th^ hemisection was processed free-floating according to standard procedures described previously (Ansorg et al., 2015). After rinsing with TBS (pH 7.4), sections were blocked in Background Sniper (Biocare Medical; for cyclin D1) or in TBSplus containing 0.2% Triton X-100, 3% or 10% donkey serum and 2% BSA (1 h at room temperature), which was also used for antibody dilution. Primary antibodies were incubated for approximately 40 hours at 4°C followed by overnight incubation in secondary antibodies with three rinsing steps in-between. To stain nuclei, DAPI (Sigma) or Hoechst33342 (Invitrogen) were added at this step. Finally, sections were rinsed three times with TBS, mounted to gelatinized slides, air-dried and coverslipped with aqueous mounting medium (Fluoromount-G, Southern Biotech or DakoCytomation Fluorescent Mounting Medium, DakoCytomation). For staining with antibodies against cyclin D1 and MCM2, we performed an epitope retrieval before the first blocking step. Therefore, sections were either steamed for 5 min in Reveal (Biocare Medical; for cyclin D1) or in sodium citrate buffer (pH6; for MCM2), followed by rapid cooling on ice water and wash-out in TBS.

Immunoperoxidase staining resembled the standard protocol described above with the following modifications: Before blocking in TBSplus, sections were treated with 1.5% H2O2 for 30 min at RT. After primary antibody incubation, sections were sequentially incubated in biotinylated secondary antibody (all raised in donkey, Dianova; 1:500), avidin-biotin-peroxidase solution (ABC Elite, Vector Laboratories, Burlingame, CA, USA) for 1 h at room temperature, followed by 3,3’-diaminobenzidine (Sigma-Aldrich) signal detection and mounting in Neo-Mount (Merck-Millipore).

For mice injected with retroviral vectors, immunostaining was performed as previously described using the methods optimized for the use in transgenic mice (Encinas and Enikolopov, 2008; Encinas et al., 2011). Animals were transcardially perfused with 30 ml of PBS followed by 30 ml of 4% (w/v) PFA in PBS, pH 7.4. Next, the brains were removed and postfixed for 3 h at room temperature in the same fixative solution, then transferred to PBS-0.2% sodium azide and kept at 4°C. Serial 70 μm-thick coronal sections were cut using a Leica VT 1200S vibratome to properly visualize the extension of the SFFV-RV infection. For immunostaining, sections were incubated with blocking and permeabilization solution (PBSplus containing 0.25% Triton-X-100 and 3% BSA) for 3 h at room temperature, and then incubated overnight with the primary antibodies (diluted in PBSplus) at 4°C. After the incubation, the primary antibody was removed and the sections were washed with PBS three times for 10 min. Next, the sections were incubated with fluorochrome-conjugated secondary antibodies diluted in PBSplus for 3 h at room temperature. After washing with PBS, the sections were mounted on gelatin coated slides and coverslipped with DakoCytomation Fluorescent Mounting Medium. The mCherry signal from SFFV-RV was detected with an antibody against DsRed.

The following antibody combinations were used: chicken anti-GFP (AvesLabs, 1:1000), goat anti-GFP (Acris, 1:300-500), mouse anti-GFAP (Millipore, 1:1000), rabbit anti-GFAP (Synaptic Systems, 1:500), goat anti-GFAP (ABCAM, 1:500), rabbit anti-S100β (DakoCytomation, 1:500), rabbit anti-cyclin D2 (Santa Cruz; 1:25), rabbit anti cyclin D1 (Thermo Scientific, 1:50), rabbit anti cyclin D1 (ABCAM, 1:500), rabbit anti-Prox1 (ReliaTech, 1:1000), guinea pig anti-DCX (Millipore; 1:500), mouse anti-MCM2 (BD Biosciences, 1:250), rabbit anti-Ki67 (Vector Laboratories; 1:750), rabbit anti-DsRed (Abcam; 1:2000), AlexaFluor-488 anti-chicken (Molecular probes; 1:500), AlexaFluor-488 anti-goat (Molecular probes; 1:500), RhX anti-guinea pig (Dianova; 1:500), RhX anti-mouse (Dianova; 1:500), RhX anti-rabbit (Dianova; 1:500), AlexaFluor647 anti-mouse (Dianova; 1:500), AlexaFluor647 anti-rabbit (Dianova; 1:500),AlexaFluor647 anti-goat (Dianova; 1:500).

## QUANTIFICATION AND STATISTICAL ANALYSIS

### Image capture and analysis

All sections were imaged on Zeiss confocal microscopes (LSM710 and LSM900; Carl Zeiss, Germany) or on a Leica SP8 confocal microscope (Leica, Wetzlar, Germany). The signal from each fluorochrome was collected sequentially, and brightness, contrast, and background were adjusted equally for the entire image using supplier’s software (Zeiss ZEN black edition and ZEN 3.0 blue edition; Leica LAS X Life Secence). The total volume of each GCL was calculated using the 10x objective of a Leica SP8 confocal microscope to completely visualize the DG in each section. The area tool of the LASX Leica software was used to measure the area of the GCL in each section. The corresponding section thickness was measured in 3 points in each slice using the z.stack setting. For the pictures of developing mouse DG, in which the DMS is illustrated, the 3DHistech panoramic digital slidescanner (Sysmex, Germany) was used.

Quantitative analysis of cell populations *in vivo* was performed by design-based (assumption free, unbiased) stereology using a modified optical fractionator sampling scheme as previously described mice (Encinas and Enikolopov, 2008; Encinas et al., 2011). For cell densities, quantifications were done maintaining the same z-stack size between conditions, corrected for the physical section thickness (measured in the GCL) and normalized to one mm^2^ SGZ. Total numbers of cells were estimated by multiplying the density with the area of the SGZ. Therefore, the length of the SGZ was determined in every slice of a single series, summed up per animal and multiplied by the section thickness (as cut), the section intervall and by two to obtain the bilateral SGZ area.

Confocal image stacks were captured along the entire rostro-caudal extend of the DG except the most caudal part (corresponding to Bregma ≤ −3.5 in adults), in which RGLs cannot be reliably identified by their morphology. Depending on the DG size along the rostro-caudal axis, one to five image stacks (512 x 512 pixels resolution) covering the SGZ/GCL/ML were random-systematically acquired per hemisection. In P0 mice, the entire DG was imaged. For quantifying NSCs, astrogliogenesis and apoptosis, a 40x/1.3 oil immersion objective was used to acquire image stacks of 100 μm x Y μm x 12.07 μm size (XYZ; 0.71 μm Z-interval). X was aligned parallel to the SGZ and Y was variably adjusted to ensure that the SGZ, GCL and ML were completely covered. For classification of NSCs, optical fields covering the SGZ/GCL were obtained with a 40x oil immersion objective. The hilus was imaged separately with the optical field size set to 10000 μm^2^. For evaluating the neuronally committed progenitor populations, 100 μm x 100 μm x 7 μm image stacks (Z-interval of 0.70 μm) covering the SGZ and GCL were captured using a 63x/1.4 oil objective.

NSCs were defined as GFP^+^GFAP^+^ cells with an apical process spanning towards the ML through at least 2/3 of the GCL (Encinas and Enikolopov, 2008). For experiments during postnatal stages prior to P10, where NSCs are still not morphologically adult-like (Brunne et al., 2013; Nicola et al., 2015), NSCs were quantified as such when they had their soma located inside the SGZ/GCL and fulfilled the following criteria: Expression of GFP and GFAP and presence of a radial process extending towards the ML. To identify proliferating NSCs, we used the marker Ki67. To classify NSC subtypes in adult mice, GFP^+^GFAP^+^ cells located in the SGZ/GCL (each GFP^+^GFAP^+^ cell in D2KO mice, approx. 50% of GFP^+^GFAP^+^ cells in WT mice) were categorized based on the expression of S100β and morphological criteria (Gebara et al., 2016; Martin-Suarez et al., 2019): α cells were defined as GFP^+^GFAP^+^S100β-cells with the soma located in the SGZ and a long radial process extensively branching at the border between GCL and ML. β cells were identified by their S100β^+^ soma located in the SGZ or GCL and a short primary process branching more proximal and rarely extending into the ML. Ω cells displayed a multibranched morphology with several primary processes extending from a S100β-soma located in the GCL. For the morphological analysis of NSCs, individual GFP^+^ NSCs whose cell bodies were located either in the SGZ or GCL were randomly selected (approx. 8-10 per animal), and imaged over their entire extend using a 63x oil immersion objective (resolution of 1024 x 1024 pixels). The obtained Z-stacks were analyzed using the 3D-Sholl analysis plugin in Fiji (http://fiji.sc/Sholl_Analysis; Ferreira et al., 2014).

Neuronally committed type 2b progenitor cells were determined by their expression of GFP, Prox1 and DCX, while type 3 neuroblasts were identified as Prox1^+^DCX^+^ cells with an apical dendrite and occasionally weak GFP expression. Astrocytes were identified by their stellate morphology and the expression of GFAP, and further distinguished into immature and mature astrocytes based on the expression or lack of GFP. To provide spatial information about potential migration routes from the germinative matrices towards the ML, quantification was conducted separately in the outer half of the GCL (oGCL), in the inner and outer half of the ML (iML, oML), and the hilus (H). The SGZ, inner GCL and subpial region were excluded from analysis, because especially until P10 these regions were too densely packed with GFP^+^GFAP^+^ cells to clearly identify astrocytes.

The number of apoptotic cells was estimated from the same Z-stacks in which D1-positive cells were phenotyped. To increase sampling accuracy in the hilus, we additionally included image stacks acquired for analysis of D2 expression and neurogenesis (approximately every second hemisection was analyzed). Apoptotic cells were identified by their densely stained and characteristically shaped nuclei, in which the aggregated chromatin appears as peripheral condensation or one solid, compact sphere (pyknosis) or fragmented into multiple spheres (karyorrhexis; Doonan and Cotter, 2008; Fig. S3).

For the experiments using SFFV-RV infections, we quantified all cells infected in the SGZ/GCL. We also quantified the number of NSCs in the infected area (region with SFFV-RV-positive cells) to obtain the percentage of NSCs infected with the SFFV-RV.

### Statistical analysis

Significance levels were assessed in Sigma Plot 14 (San Jose, CA, USA). Data were first tested for normality. If given, either a two-tailed Student’s T-test or a Welch’s T-test (in case of no homoscedasticity) were applied to compare a single dependent variable between two groups, otherwise we used a Mann-Whitney-U test. For comparing a single dependent variable between more than two groups, we applied a One-way ANOVA, while 2-way ANOVAs were applied to assess the effect of two independent variables on one dependent variable. A One-way repeated measures (RM) ANOVA was used for comparing several dependent variables in one group. To examine the effect of two independent variables, of which one been has been measured repeatedly, we applied a 2-RM ANOVA. All ANOVAs were followed by Tukey’s post-hoc tests to determine specific differences. A *p* < 0.05 was considered as statistically significant. All data represent mean ± SEM.

## TABLE FOR AUTHOR TO COMPLETE

Please upload the completed table as a separate document. **<u>Please do not add subheadings to the key resources table.</u>** If you wish to make an entry that does not fall into one of the subheadings below, please contact your handling editor. **<u>Any subheadings not relevant to your study can be skipped.</u>** (**NOTE:** For authors publishing in Cell Genomics, Cell Reports Medicine, Current Biology, and Med, please note that references within the KRT should be in numbered style rather than Harvard.)

### Key resources table (blue = examples, need to be deleted)

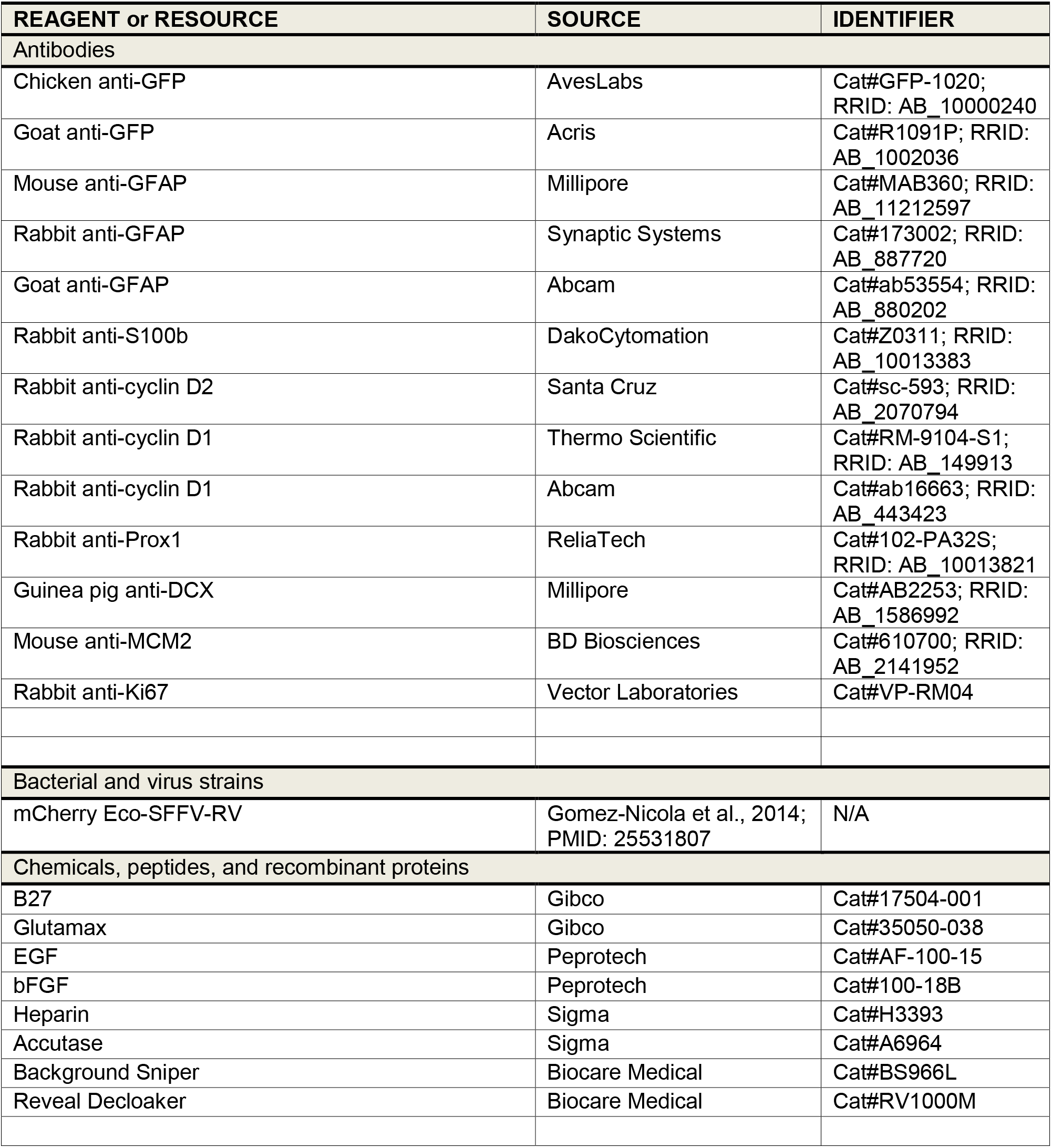
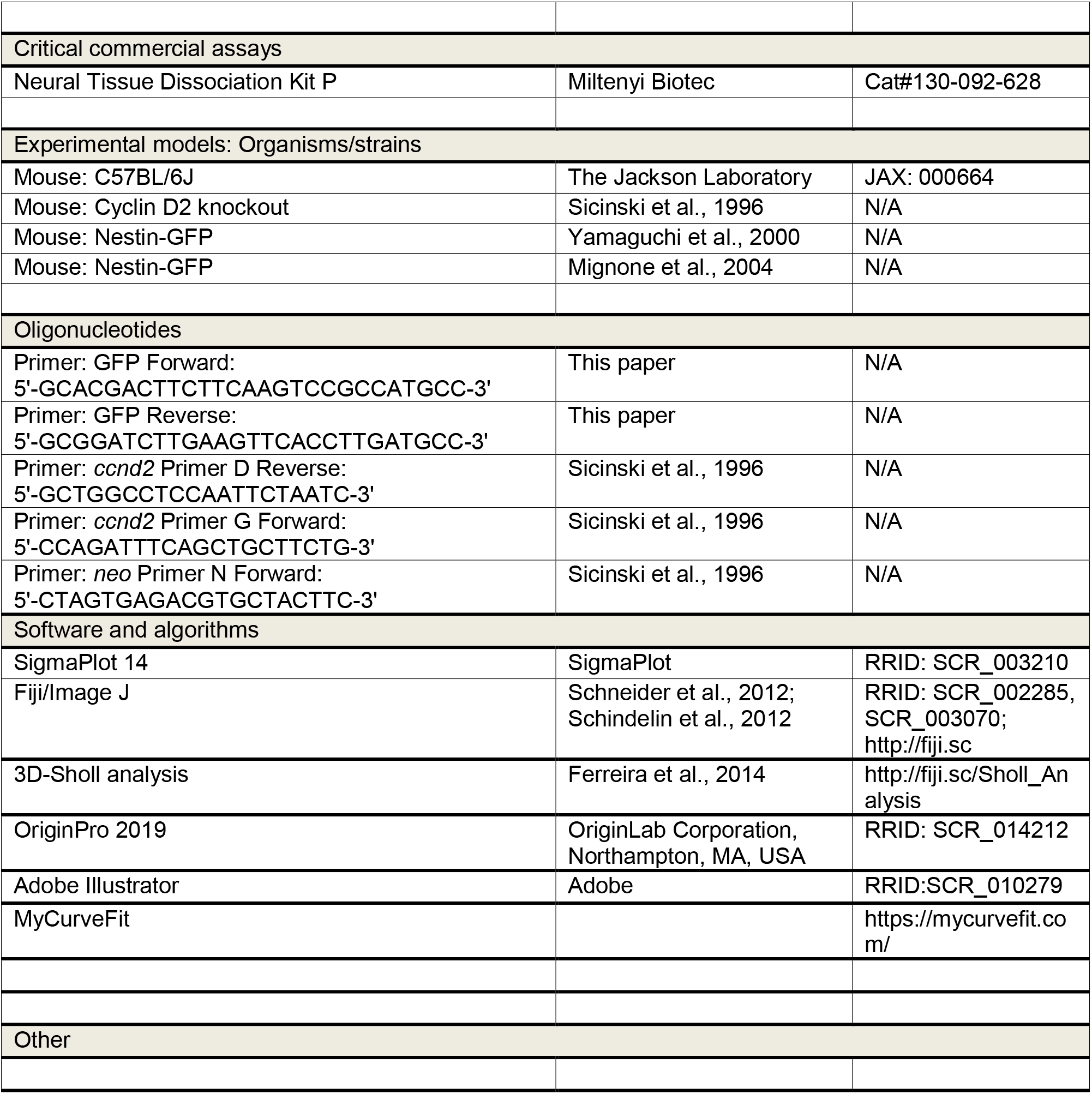

