## Supplementary figures and images for "Generation of adult hippocampal neural stem cells occurs in the early postnatal dentate gyrus and depends on cyclin D2"

### Supplemental Figure 1

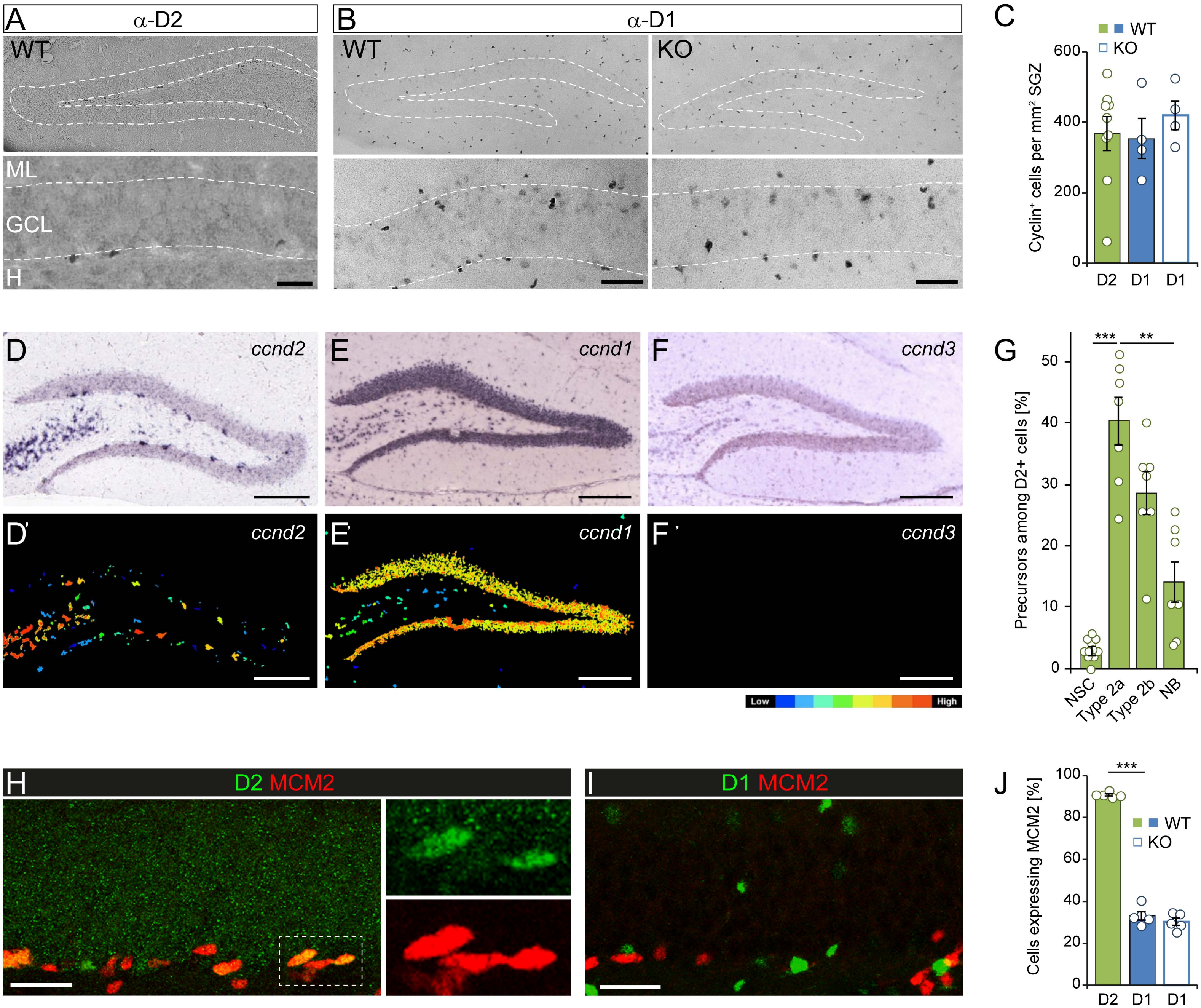

### Supplemental Figure 2

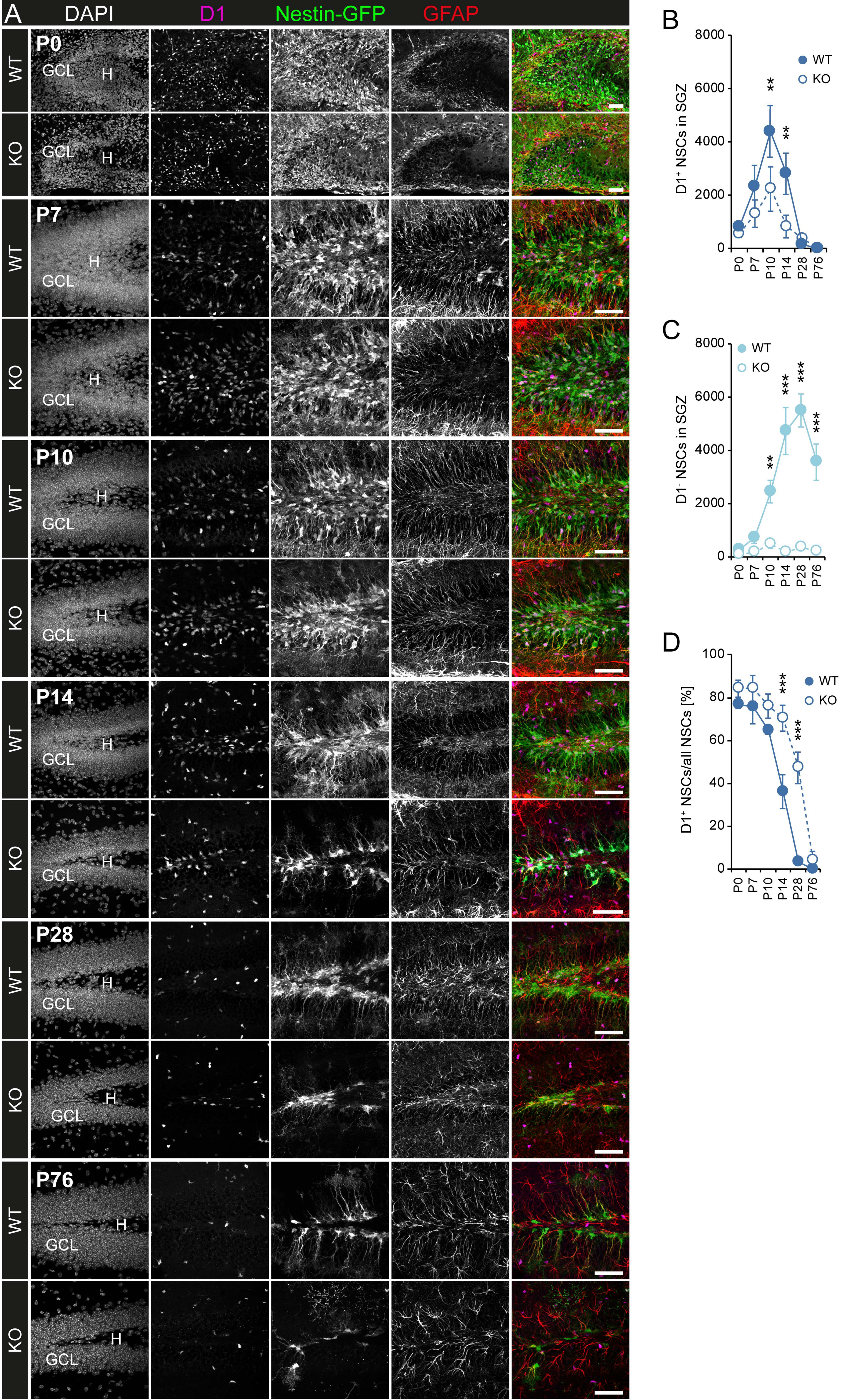

### Supplemental Figure 3

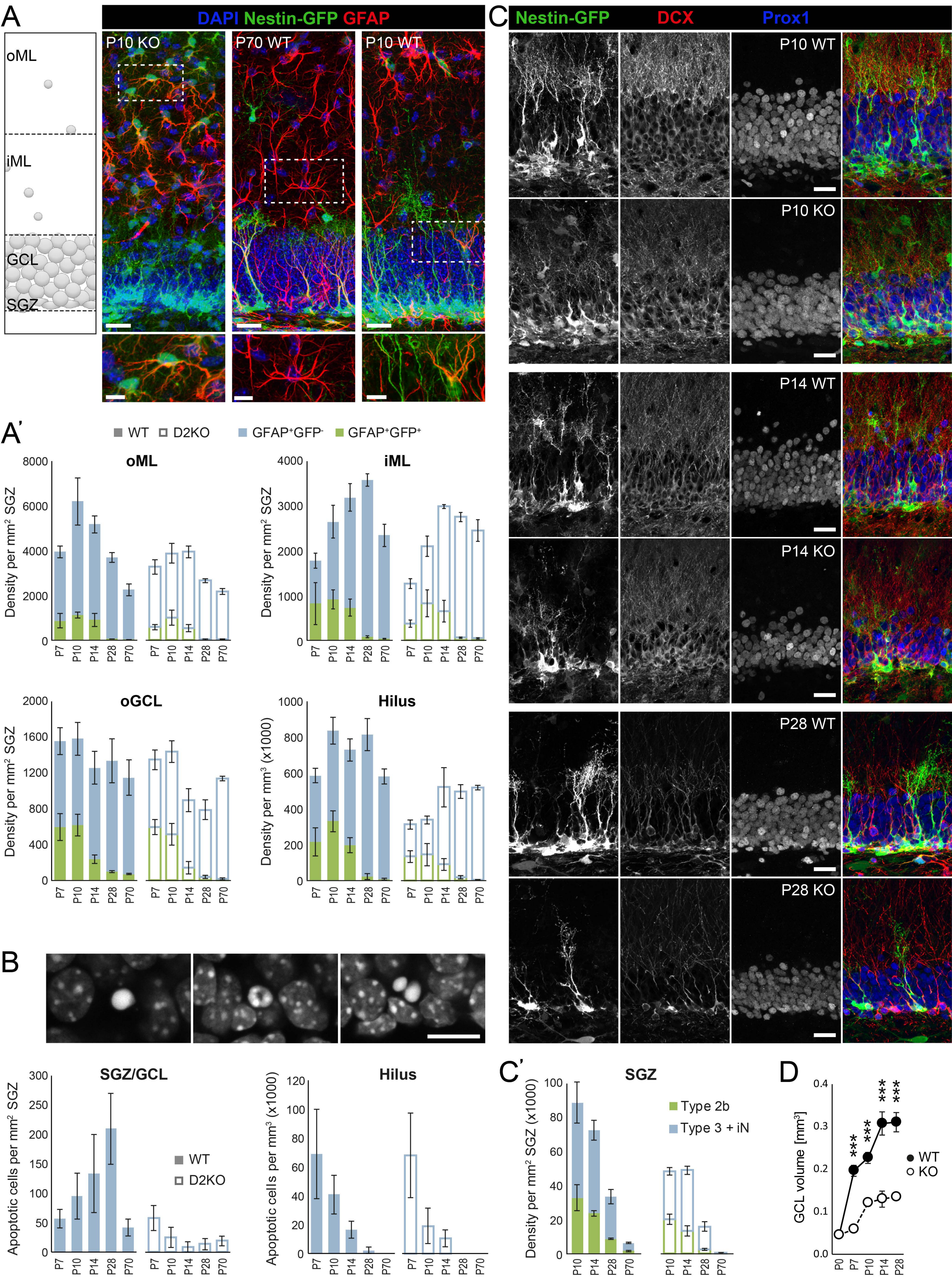

### Supplemental Figure 4

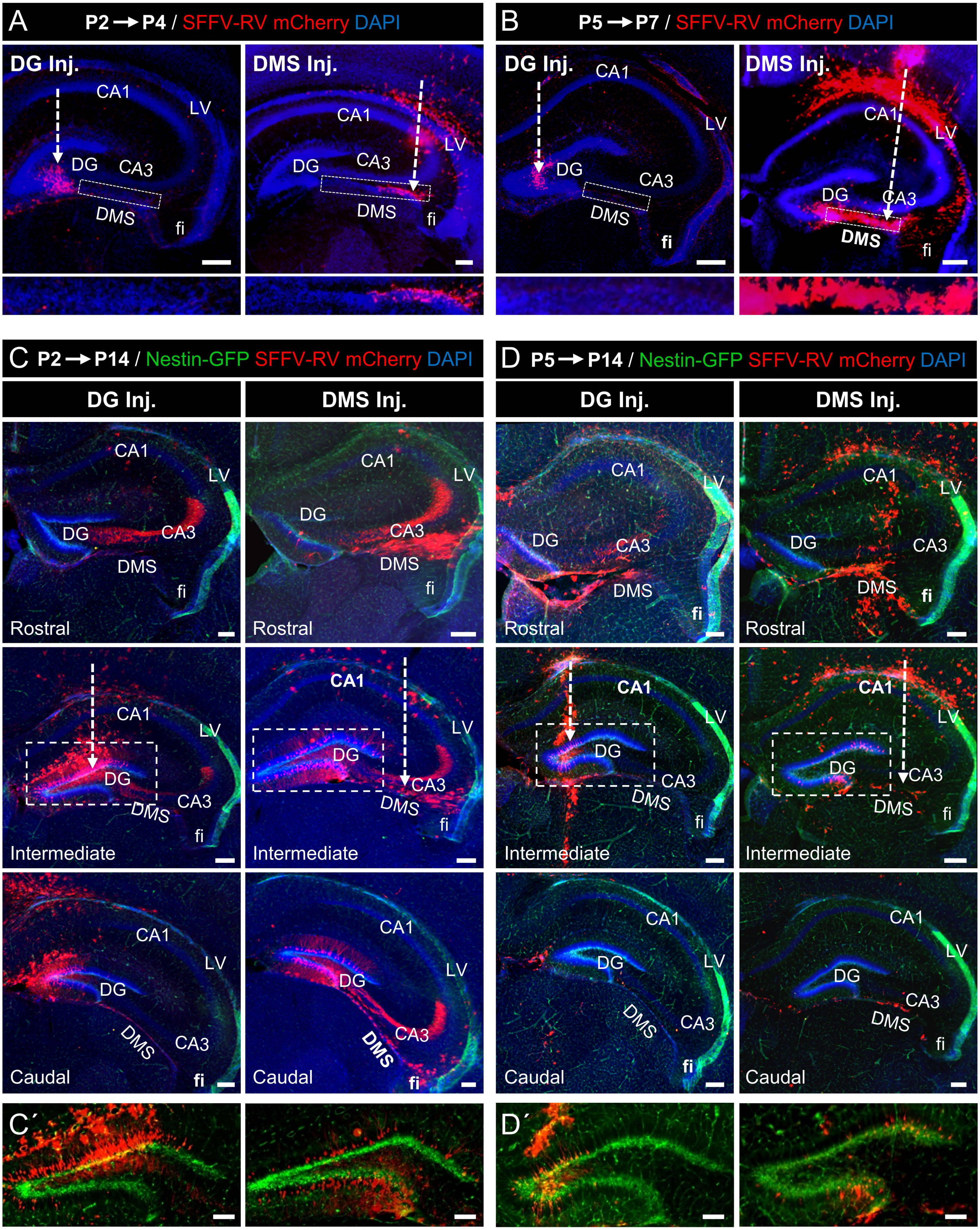
